# Optogenetic clustering and membrane translocation of the BcLOV4 photoreceptor

**DOI:** 10.1101/2022.12.12.520131

**Authors:** Ayush Aditya Pal, William Benman, Thomas R. Mumford, Brian Y. Chow, Lukasz J. Bugaj

## Abstract

Optogenetic clustering is a versatile method to control protein activity in living cells, tissues, and organisms. Here we show that the BcLOV4 photoreceptor both clusters and translocates to the plasma membrane in response to blue light, representing a new class of light-dependent behavior. We demonstrate that dual translocation and clustering can be harnessed for novel single-component optogenetic tools, including for activation of the entire family of epidermal growth factor receptor (ErbB1-4) tyrosine kinases. We further find that clustering and membrane translocation are causally linked. Stronger clustering increased the magnitude of translocation and downstream signaling, increased sensitivity to light by ~3-4-fold, and decreased the expression levels needed for strong signal activation. Thus light-induced clustering of BcLOV4 provides a strategy to generate a new class of optogenetic tools and to enhance existing ones.

## Introduction

Optogenetics enables optical control of proteins by coupling them to naturally evolved photoreceptors. These photoreceptors can undergo one of a handful of inducible behaviors, including conformational changes^1,2^, homo/hetero-dimerization^3–8^, clustering^9,10^, and membrane translocation^11,12^, each of which has been leveraged to control numerous aspects of cell physiology. In some cases, a photoreceptor can possess multiple such functions. One example is *Arabidopsis* Cry2, which, in addition to heterodimerizing with CIB1, also forms light-induced clusters^8–10,13^.

BcLOV4 is a photoreceptor that translocates from the cytosol to the plasma membrane under blue light^11^. BcLOV4 translocation has been leveraged for multiple probes of cell signaling, including of Rho GTPases, Ras, and PI3K signaling, and works across organisms including yeast, flies, zebrafish, and mammals^11,14–16^. While microscopy showed that, in cells, stimulated BcLOV translocates to the plasma membrane, experiments with purified protein found that BcLOV can also undergo light-induced aggregation^11^. However, clustering was only observed in the absence of lipid membranes. Within water-in-oil emulsions, light-stimulated BcLOV localized to the membrane and did not visibly aggregate, although aggregation was still observed in the center of large emulsion droplets where the diffusive distance to the membrane was largest^11^. Collectively, these results suggested that BcLOV4 clustering and membrane association may be mutually exclusive. However, whether BcLOV4 forms clusters at the membrane, and whether clustering plays a role in BcLOV translocation, has not been formally tested.

The inability to observe BcLOV4 clustering in cells could be explained if membrane-associated clusters were sufficiently small. Small clusters will not appear punctate under conventional fluorescence imaging due to measurement limitations including the diffraction limit of light and a low signal-to-noise of the aggregated fluorophore against the fluorescence background^17^. A similar effect can be observed with the *bona fide* clustering module Cry2, which clusters in response to blue light, but whose clusters can only be observed above an expression threshold^9,17–20^. Recently our group developed the CluMPS reporter to indicate the presence of protein clusters as small as trimers^17^. When applied to Cry2, CluMPS revealed the presence of small Cry2 clusters at all expression levels, including at low expression levels where clusters could not be otherwise observed^17^.

Clarifying the existence of BcLOV4 clustering in cells would be impactful for several reasons. Optogenetic clustering of Cry2 has been a powerful approach, for example in studies of cell signaling^9^,^21^, stem cell differentiation^22^,^23^, neurodegenerative aggregation^24^, and protein phase separation^25^. However, Cry2 remains the only photoreceptor whose light-induced clustering has been used for optogenetic control. Additional such methods would expand the applications towards which optogenetic clustering could be applied. Further, understanding the molecular details of BcLOV activation may yield insights to understand and mitigate the unique temperature-sensitivity of BcLOV4, which spontaneously self-inactivates within ~ 1 hr of strong light stimulation above ~30 °C^16^.

In this work, we find that BcLOV4 is a multifunctional protein that concurrently clusters and translocates to the membrane under light stimulation. We leverage this multifunctionality to generate new tools for the activation of the epidermal growth factor receptor (EGFR) kinase, which could not be activated by membrane translocation alone. We further apply this same strategy to activate the entire ErbB receptor family in a modular manner and find receptor-specific signal dynamics. Surprisingly, in contrast to previous evidence that clustering and translocation are antagonistic, we find that clustering potentiates BcLOV4 membrane translocation, sensitizes stimulation to lower levels of light, and diminishes temperature-dependent inactivation. Our work thus uncovers new features of BcLOV4 stimulation and provides a platform to engineer a unique class of optogenetic tools and to enhance existing ones.

## Results

To examine whether BcLOV4 formed light-induced clusters at the membrane, we transiently transfected BcLOV-GFP in HEK 293T cells and observed its distribution after blue light stimulation (**Figure 1A**). As reported before^11,14–16,26^, fluorescence appeared mostly uniform at the membrane (**Figure 1B)**. However, we noticed that in cells with high expression, light-induced fluorescent puncta could be observed at the membrane (**Figure S1A**). Since cluster size depends on concentration, we reasoned that smaller clusters may be forming in low-expressing cells as well, but at a submicroscopic scale. To test this possibility, we repeated our imaging experiment in the presence of a CluMPS reporter. A CluMPS reporter amplifies small protein clusters through multivalent interactions that generate large fluorescent condensates in the presence of a clustered target ^17^ (**Figure 1C**). In cells that co-expressed BcLOV-GFP with a CluMPS reporter of GFP clustering (LaG17-CluMPS), light stimulation rapidly triggered both membrane-association and condensate formation regardless of BcLOV4 expression level (**Figure 1D, Figure S1B, Supplementary Movie 1**). Notably, CluMPS did not produce reporter condensates in response to membrane recruitment of GFP through optogenetic heterodimerization (iLid and sspB-GFP^5^), suggesting that clustering and CluMPS activation was a not a general property of membrane translocation (**Figure S1C**).

**Figure 1:**
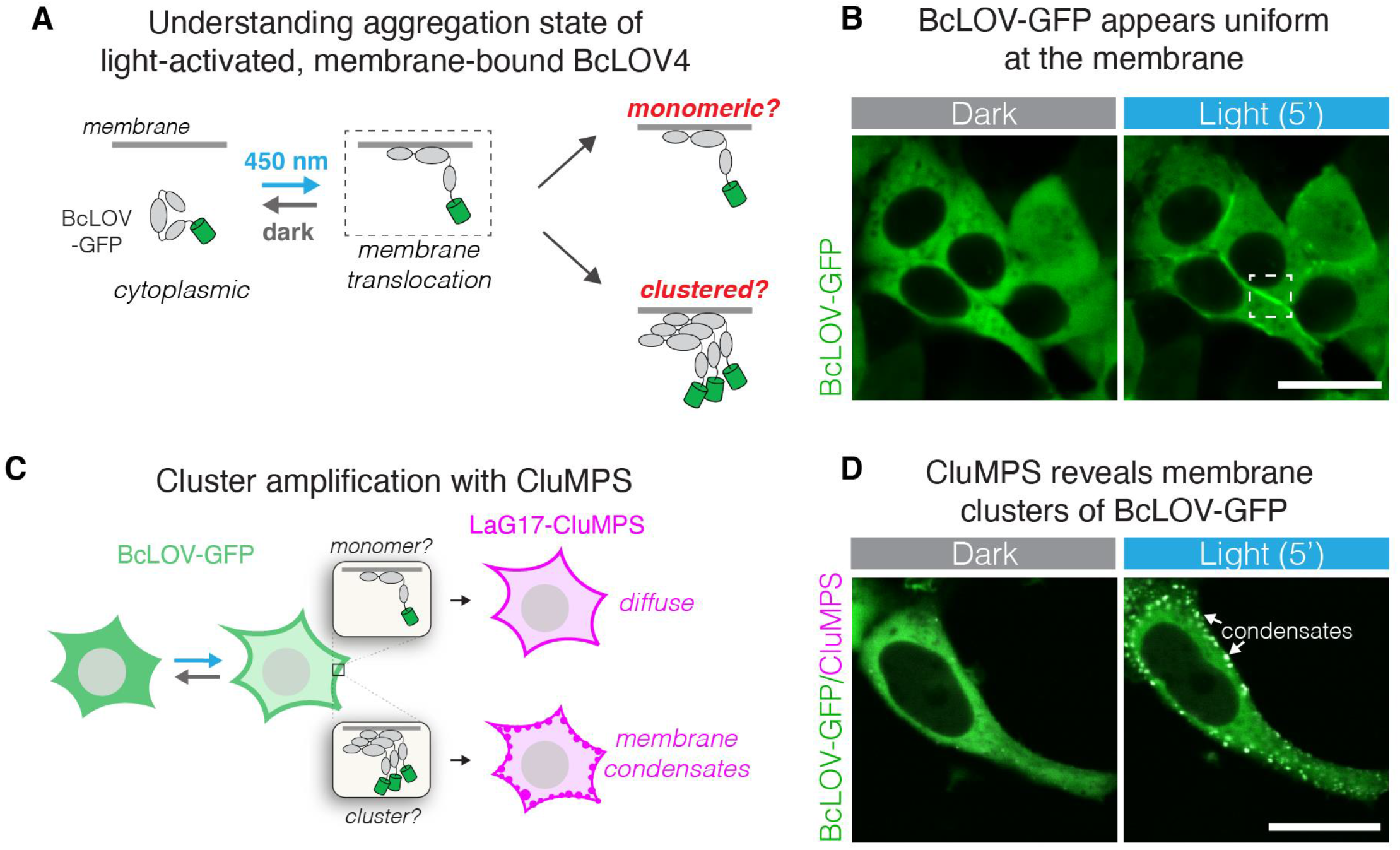
BcLOV4 forms light-induced clusters at the membrane. **A)** BcLOV-GFP translocates to the plasma membrane when stimulated with blue light. However, it is unknown whether it forms clusters at the membrane. **B)** Representative image of membrane recruitment of BcLOV-GFP upon blue-light stimulation in HEK 293T cells. Dashed box shows membrane localization as depicted in (A). Scale bar = 20 μm. **C)** The CluMPS reporter for GFP clustering (LaG17-CluMPS) was co-expressed with BcLOV to amplify and visualize potential submicroscopic membrane-associated clusters of BcLOV-GFP. **D)** Representative images of membrane recruitment of BcLOV-GFP in the presence of LaG17-CluMPS. CluMPS amplifies and visualizes membrane-associated BcLOV condensates in the light. Scale bar = 20 μm (See **Supplementary Movie 1)**. See **Supplementary Figure 1** for additional controls and **Supplementary Table 1** for details of optogenetic illumination parameters.

The unique ability of BcLOV4 to both translocate and cluster in response to blue light carries the potential for new types of optogenetic tools. The optogenetic clustering protein Cry2 has been applied to cluster and activate receptor tyrosine kinases (RTKs)^21^,^27–30^. However, these tools required either constitutive anchoring to the plasma membrane, which could raise basal signaling levels^27–29^,^31^, or required a separate interaction partner anchored at the membrane, which can necessitate stoichiometric tuning between the two components for optimal function^31^,^32^. We reasoned that BcLOV4 could implement a simpler, single-component variant of such tools, without the need for membrane anchoring.

We first sought to stimulate EGFR (ErbB1), a receptor important for cell growth and survival that is commonly misregulated in human cancers. We fused BcLOV-mCh to the N-terminus of the EGFR intracellular domain (BcLOV-EGFR, **Figure 2A**). To assay activation, we observed activity of the downstream Erk kinase using the ErkKTR reporter, a fluorescent probe that translocates from the nucleus to the cytoplasm upon Erk activation (**Figure 2B**)^33^. Within seconds after light stimulation, BcLOV-EGFR translocated to the membrane, and within minutes, ErkKTR-miRFP moved from the nucleus to the cytoplasm, indicating Erk activation (**Figure 2C, Supplementary Movie 2**). BcLOV-EGFR signaling could be stimulated and inactivated over multiple cycles (**Figure 2D**). Immunofluorescence staining for phospho-Erk (ppErk) confirmed light induced Erk activation and also showed a lack of basal pathway activation in transfected but unstimulated (dark state) cells (**Figure S2**). Notably, ErkKTR activation and ppErk elevation were not observed in cells where the EGFR domain was recruited to the membrane through 1:1 heterodimerization of the iLid/sspB system (**Figure S3**) confirming that 1) both membrane translocation and clustering are required for activation of the EGFR intracellular domain, and 2) BcLOV clustering at the membrane can be leveraged for novel optogenetic signaling tools.

**Figure 2:**
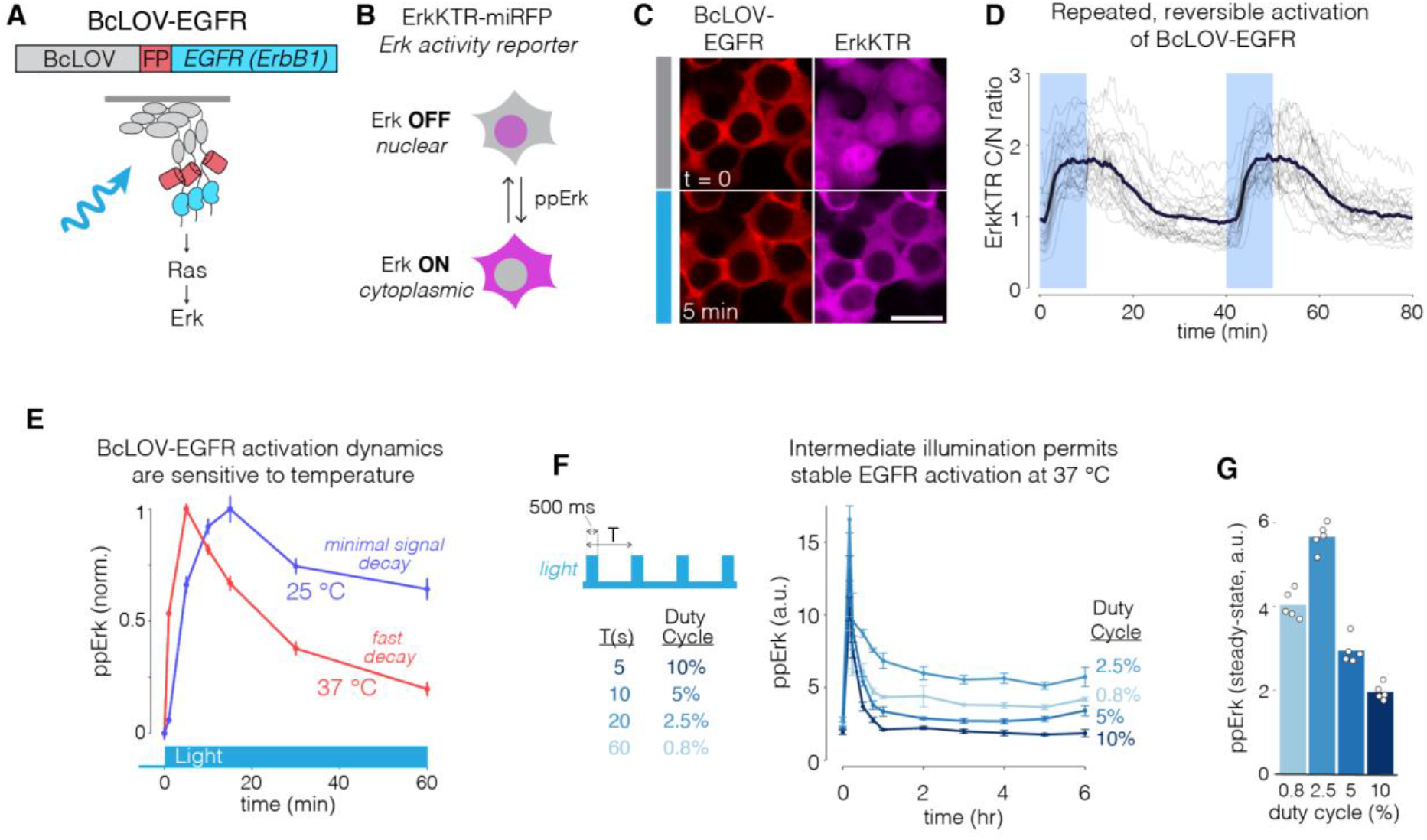
Membrane translocation and clustering allows activation of EGFR. **A)** The intracellular domain of EGFR was fused to BcLOV-mCherry to generate BcLOV-EGFR. Light-induced BcLOV-EGFR activity was assessed by measuring activity of downstream Ras-Erk signaling. **B)** ErkKTR is a fluorescent biosensor of Erk activity. ErkKTR is nuclear when Erk is off and is cytoplasmic when Erk is on. **C)** HEK 293T cells that coexpress BcLOV-EGFR and ErkKTR-miRFP show Erk activation upon light stimulation. Scale bar = 20 μm (See **Supplementary Movie 2**). **D)** Erk activity could be stimulated reversibly over multiple cycles. Grey traces represent mean ErkKTR cytoplasmic/nuclear ratios of individual cells (n = 25). Black trace represents mean of these traces. **E)** BcLOV-EGFR signal kinetics depend on temperature. Erk signal can be stably maintained with light at 25 °C but decays more rapidly 37 °C. Data represent mean ± SEM of four replicates, with each replicate representing the mean of ~1000-4000 cells. **F)** BcLOV-EGFR activity can be stably maintained at 37 °C at intermediate light doses. Cells were stimulated with pulse trains of light of variable duty cycles. **(G)** Maximal steady-state Erk levels were achieved at 2.5% duty cycle (500 ms ON every 20 s). Data in (G) represents the mean ± SEM of four replicates, with each replicate representing the mean of ~300-1700 cells. Datapoints in (G) are the mean steady-state (from 2 hr to 6 hr) ppErk levels shown in (**F**). See **Supplementary Table 1** for details of optogenetic stimulation for all experiments. a.u., arbitrary units.

BcLOV4 membrane translocation dynamics depend on both light and temperature^16^. Although BcLOV binds the membrane at all temperatures, it then spontaneously dissociates within ~ 1 hr at a rate that depends on both temperature and light exposure. BcLOV-based signaling probes also showed self-inactivation in cells cultured at 37 °C^16^. We observed similar temperature-dependent behavior for BcLOV-EGFR: Erk phosphorylation could be stably maintained at 25 °C, but decayed more rapidly within ~ 1 hr of strong light stimulation at 37 °C (**Figure 2E**). However, in contrast to complete inactivation of BcLOV-SOS_cat_, we found that BcLOV-EGFR could sustain intermediate levels of pathway activation (> 6 hr) at mammalian temperatures by using an intermediate doses of stimulating light (**Figure 2F,G**). Such intermediate doses sustain signaling presumably because they stimulate a sufficient amount of BcLOV for EGFR activation, but also a small enough amount such that only a small fraction of BcLOV total undergoes inactivation, leaving a large reservoir of activatible BcLOV to maintain signal activity under sustained stimulation.

EGFR is a member of the ErbB receptor family, whose members (ErbB1-4) play important roles in development as well as cancer^34^. However, it is currently challenging to study the specific activity of each ErbB family member in isolation for several reasons. First, receptor ligands can activate multiple family members. Second, ErbB2 has no known ligand. Third, the ErbB family members can heterodimerize with each other upon ligand activation. Chemical and optical probes have been developed to overcome this challenge for EGFR and ErbB2^30,35,36^, although individual methods that can stimulate each member of the ErbB receptor family have not been reported. We thus asked whether BcLOV clustering could be used to stimulate ErbB2-4 in the same manner as for EGFR. We fused the intracellular domains of each ErbB receptor to the C-terminus of BcLOV-mCh and observed membrane translocation under light stimulation (**Figure 3A)**. Each fusion rapidly localized to the plasma membrane after stimulation with blue light (**Figure 3B)**. Intriguingly, the magnitude of translocation differed between fusions. ErbB1(EGFR) and ErbB2(Her2) showed weak-to-moderate translocation, with apparent uniformity of fluorescence at the membrane. In striking contrast, ErbB3 showed the strongest translocation, even stronger than the original BcLOV-mCh fusion, and showed obvious clusters at the membrane. ErbB4 showed strong membrane translocation and moderate membrane clustering, lower than ErbB3 but more than ErbB1/2. Notably, ErbB3 and ErbB4 fusions on occasion formed cytoplasmic condensates in the dark, and these condensates would dissolve in favor of membrane translocation after light stimulation (**Figure 3B, Supplementary Movie 3**).

**Figure 3:**
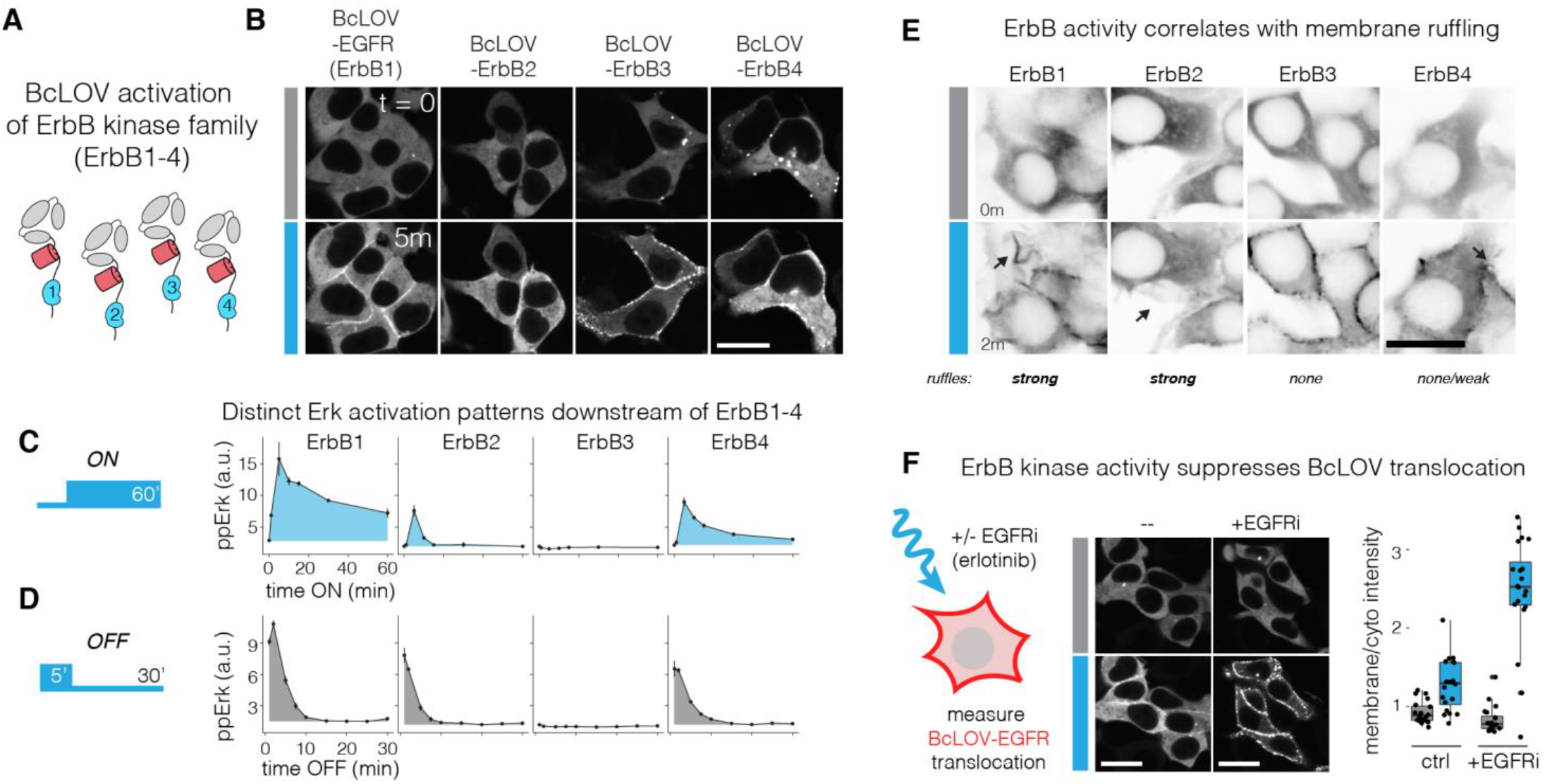
BcLOV4 clustering at the membrane allows for modular activation of the entire ErbB receptor family. **A)** The intracellular domains of ErbB1-4 were fused to the C-terminus of BcLOV-mCherry. **B)** Membrane translocation of BcLOV-ErbB1-4 in response to blue light. Scale bar = 20 μm. See also **Supplementary Movie 3**. **C,D)** Erk activation dynamics downstream of BcLOV-ErbB1-4 in response to ON and OFF steps of blue light. See also **Supplementary Figure 4**. Data represent mean ± SEM of two replicates, with each replicate representing the mean of ~500-2400 cells. **E)** Membrane ruffling (black arrows) downstream of stimulation of BcLOV-ErbB1-4, indicative of RTK stimulation. Ruffling is strongest for ErbB1 and ErbB2, less for ErbB4, and absent for ErbB3 activation. Scale bar = 20 μm. See also **Supplementary Movie 4**. **F)** Kinase activity suppresses BcLOV-EGFR membrane translocation. Translocation was observed under light stimulation in the presence or absence of 1 μM EGFR inhibitor erlotinib (EGFRi). EGFRi promoted stronger translocation. Scale bar = 20 μm. Quantification (right) shows ratios of mean membrane and cytoplasmic fluorescence of 25 single cells.

We next asked whether translocation of the BcLOV-ErbB fusions activated downstream signaling by measuring downstream Erk phosphorylation. Each receptor fusion elicited distinct Erk activation dynamics (**Figure 3C**). In response to sustained stimulation, ppErk activation was strongest for ErbB1, whereas ErbB2 and ErbB4 showed weaker signaling despite equivalent expression levels (**Figure S4**). ErbB1 and ErbB4 also showed sustained signal above baseline, whereas ErbB2 signaled with a transient pulse and rapidly returned to baseline (**Figure 3C, Figure S4**). ErbB3, by contrast, showed no Erk phosphorylation, in line with the fact that ErbB3 is a pseudokinase and lacks enzymatic activity^37–39^. We found no major differences in the OFF-kinetics between each tool, as measured after 5 minutes of light stimulation and subsequent light removal (**Figure 3D)**. The half-life of signal decay was ~ 5 minutes for Erb1,2 and 4, with complete loss of signal by 15 minutes. We further confirmed signal activation using the ErkKTR reporter (**Figure S5**) and through the observation of membrane ruffling, indicative of PI3K/Rac1 activation downstream of receptor activation (**Figure 3E, Supplementary Movie 4**). Collectively, our results show that BcLOV can be applied in a modular fashion, with no further optimization, to generate optogenetic tools for each of the ErbB family members.

Successful control of the ErbB receptor family led us to ask whether the modularity of BcLOV would extend to other families of RTK signals. We generated fusions of BcLOV to the intracellular domain of two other RTKs, fibroblast growth factor 1 (FGFR) and platelet derived growth factor (PDGFRβ). Both constructs could stimulate the ErkKTR reporter in HEK 293T cells (**Figure S6)**. However, for BcLOV-FGFR1, high basal ppErk and Erk activity were observed and were strongly correlated with expression levels of the fusion, such that optimal switching of Erk (OFF in dark, ON in light) could only be achieved in low-expressing cells (**Figure S6A**). FGFR1 stimulation was substantially weaker when recruited to the membrane through iLid/sspB heterodimerization, although Erk activity could still be stimulated in a fraction of cells (**Figure S6B**). BcLOV-PDGFR showed no elevation of basal ppErk in low/medium-expressing cells, and ppErk induction required medium-high expression (**Figure S6C**). Clustering was required for PDGFR activation, as membrane recruitment with iLid/sspB heterodimerization did not increase ppErk levels (**Figure S6D**). In sum, we find that BcLOV can regulate diverse RTKs in a modular manner, although receptor-specific expression, host-cell dependencies, and molecular context will dictate optimization for each individual RTK, as observed previously^28,29,35,36^.

Among the BcLOV-ErbB probes, the BcLOV-ErbB3 fusion showed the strongest translocation to the membrane despite a lack of downstream signaling (**Figure 3B**). Because ErbB3 is the only pseudokinase among the ErbB family, this result suggested that kinase activity may suppress membrane translocation of the optogenetic probe. To test this hypothesis, we measured light-induced translocation of BcLOV-EGFR, in the presence or absence of the EGFR inhibitor erlotinib (1 μM, EGFRi). Without EGFRi, BcLOV-EGFR showed only moderate translocation, as before (**Figure 3F**). However, in the presence of EGFRi (*i.e*. in the absence of EGFR kinase activity), translocation was markedly stronger, with large clusters appearing at the membrane similar to those observed for BcLOV-ErbB3 (**Figure 3F**). These results confirm that ErbB kinase activity suppresses translocation. Although the specific mechanism of suppression is not clear, we found that EGFRi treatment also enhanced EGFR recruitment using the heterodimeric iLid/sspB system (**Figure S7**). Thus kinase-dependent suppression of translocation is not specific to BcLOV4 and may be a more general feature of inducible membrane recruitment.

*In vitro* experiments with BcLOV previously suggested that its clustering and membrane association may be mutually antagonistic^11^. However, throughout our study we observed a correlation between increased clustering and stronger membrane binding (**Figure 1D**, **3B, 3F**). We thus directly tested the role of clustering on membrane association by strengthening the clustering potential of BcLOV with intrinsically disordered regions (IDRs), which have previously been used to potentiate the clustering strength of Cry2^25^ (**Figure 4A**). We tested two IDRs: the FUS low complexity domain (FUS(LC)) and the RGG domain from LAF-1, two well-characterized domains that have been used to engineer protein phase separation ^25,40,41^. Both IDR fusions dramatically enhanced optogenetic membrane translocation of BcLOV-mCh, supporting a causal positive role for clustering on translocation (**Figure 4B, Supplementary Movie 5**). Notably, both IDR fusions retained clear membrane localization of BcLOV even after two hours of stimulation at 37 °C, whereas wt BcLOV4 translocation decayed back to unstimulated levels, as observed previously^16^ (**Figure 4C**). Thus the modulation of cluster properties can tune the amplitude and temperature-dependent dynamics of BcLOV stimulation (**Figure 4D**).

**Figure 4:**
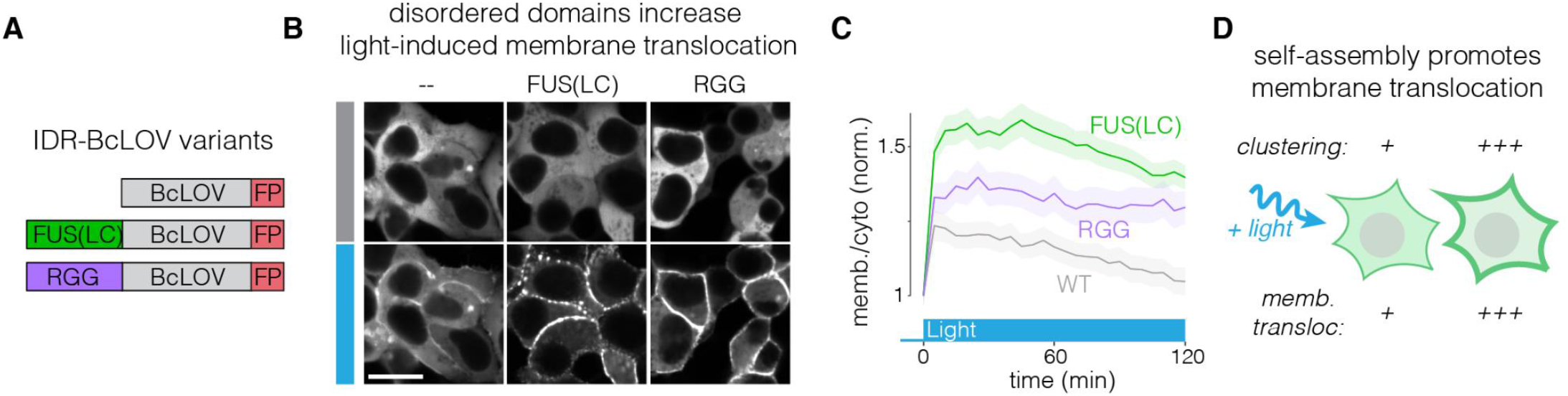
Enhanced clustering strengthens light-induced membrane binding of BcLOV4. **A)** The IDRs FUS(LC) and RGG were fused to BcLOV-mCh to test whether increasing BcLOV4 clustering strength could tune the magnitude of membrane binding. **B)** Both IDR-fused variants of BcLOV-mCh showed dramatic enhancement in membrane translocation. Scale bar = 20 μm. See also **Supplementary Movie 5**. **C)** IDR-BcLOV fusions maintained strong membrane localization even after 2 hours of stimulation at 37°C, whereas wt BcLOV-mCh decays back to unstimulated levels. Data represent mean BcLOV4 membrane/cytoplasmic ratios of ~350-750 cells. Ribbons = 95% CI (see **Methods** section for quantification details). **D)** The clustering strength of BcLOV can tune its ability to translocate to the membrane in response to light stimulation.

The ability of IDRs to tune membrane association suggested that IDRs might also tune activation of BcLOV-based tools. To test this, we first observed the effects of IDRs on activation of BcLOV-EGFR (**Figure 5A**). Both wt and FUS- or RGG-fused BcLOV-EGFR expressed well and stimulated the ErkKTR reporter in HEK 293T cells (**Figure 5B**). To quantify the effects of the IDRs, we performed a dose-response experiment to examine ppErk levels in response to a range of blue light intensities. FUS-BcLOV-EGFR drove a 30% higher maximum ppErk activation relative to wt BcLOV-EGFR, and showed a 2-fold increased sensitivity to light (1.2 vs 2.5 mW/cm^2^ to reach half-max amplitude of wt, **Figure 5C**). Interestingly, RGG-fused BcLOV-EGFR showed no benefit over the wt variant (**Figure 5C**). The divergent effects of FUS and RGG IDRs suggest that IDRs may not all work interchangeably and that enhancements in membrane translocation may not necessarily translate to enhancement of BcLOV-based optogenetic probes. Nevertheless, the increased sensitivity and responsiveness of FUS-BcLOV-EGFR permitted higher levels of ppErk during both short- and long-term stimulation as compared to the wt BcLOV-EGFR probe (**Figure 5D**). As before, maximal steady-state ppErk stimulation was observed at low-intermediate light patterns (2.5% duty cycle of 160 mW/cm^2^ blue light) (**Figure 5E**). Importantly, FUS also increased the efficiency of the probe, where equivalent signal strength could be achieved at lower expression levels compared to unmodified BcLOV-EGFR. (**Figure 5F**). Thus, amplification of BcLOV clustering and membrane translocation can generate optogenetic probes with higher sensitivity and signal strength, stronger sustained signaling, and lower requirements for probe expression levels.

**Figure 5:**
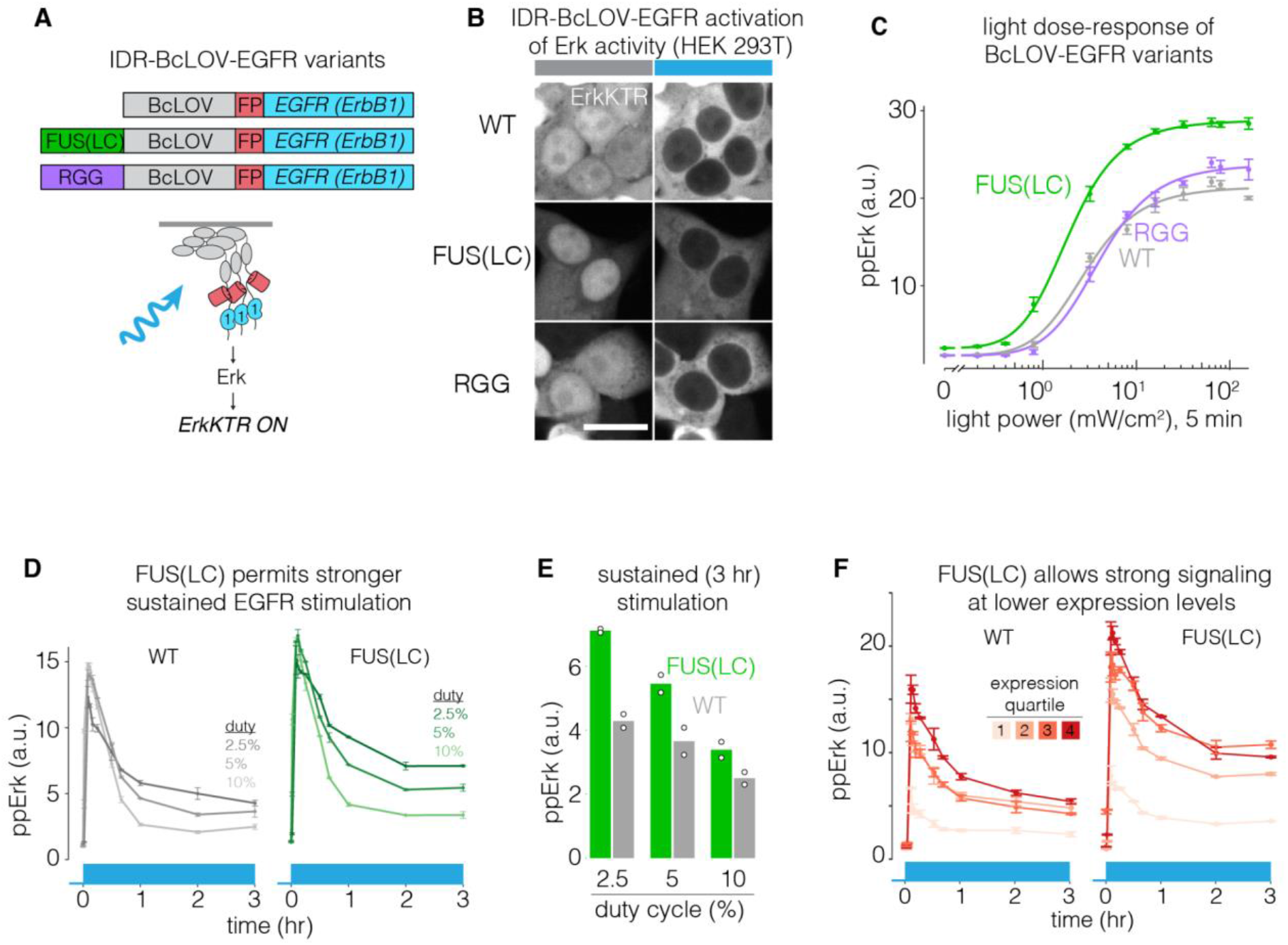
IDRs enhance sensitivity and strength of BcLOV-EGFR. **A)** IDR-fused variants of BcLOV-EGFR. **B)** IDR-BcLOV-EGFR variants stimulated ErkKTR-miRFP under blue light in HEK 293T cells. Scale bar = 20 μm. **C)** Dose-response of light intensity on ppErk after 5 min of constant illumination at the indicated light dosages. Data represent the mean ± SEM of four replicates, each representing the mean from ~2000-4000 cells. **D)** Comparison of sustained stimulation of wt or FUS-fused BcLOV-EGFR at variable duty cycles of stimulation. See **Supplementary Table 1** for details of optogenetic stimulation parameters. **E**) Steady-state levels of ppErk after 3 hours of stimulation. FUS-BcLOV-mCh allowed stronger steady state levels of ppErk at all duty cycles tested. For (D), data represent mean ± SEM of two replicates, each representing the mean of ~2000-5000 cells. Datapoints in (E) are the mean steady-state ppErk levels at 3 hr shown in (E). **F)** FUS(LC) decreases the concentration of the BcLOV probe required to achieve a given signaling level. Data points represent mean ± SEM of two replicates, with each replicate representing the mean of 400-1500 cells per expression quartile.

To determine whether benefits of increased would extend to other BcLOV-based tools, we tested the effects of IDRs on BcLOV-SOS_cat_, which stimulates Ras-Erk signaling, and which also self-inactivates at 37 °C ^16^ (**Figure 6A**). Both IDR-BcLOV-SOS_cat_ variants stimulated ErkKTR in NIH 3T3 cells (**Figure 6B**). Whereas Erk activity began to decay shortly after its rapid activation by wt BcLOV-SOS_cat_, activity was more sustained and showed slower decay when driven by either IDR-BcLOV-SOS_cat_ variant (**Figure 6C)**. Dose-response experiments showed that, while all variants reached equivalent maximal ppErk levels, both IDR variants were 3-4X more sensitive to light than wt BcLOV-SOS_cat_ (intensity for half-max activation: wt: 20 mW/cm^2^, FUS: 5.1 mW/cm^2^, RGG: 6.7 mW/cm^2^, **Figure 6D**). When illuminated at light levels that gave equivalent max ppErk response, the IDR variants yielded more sustained and integrated ppErk signal over 1 hr of constant stimulation (**Figure 6E**). We also compared signaling in response to a strong but pulsatile light input, a commonly used stimulation pattern that minimizes phototoxicity^42^ (**Figure 6F)**. Here, both IDR variants achieved > 2-fold higher maximal signal and more sustained activity compared to wt BcLOV-SOS_cat_.

**Figure 6:**
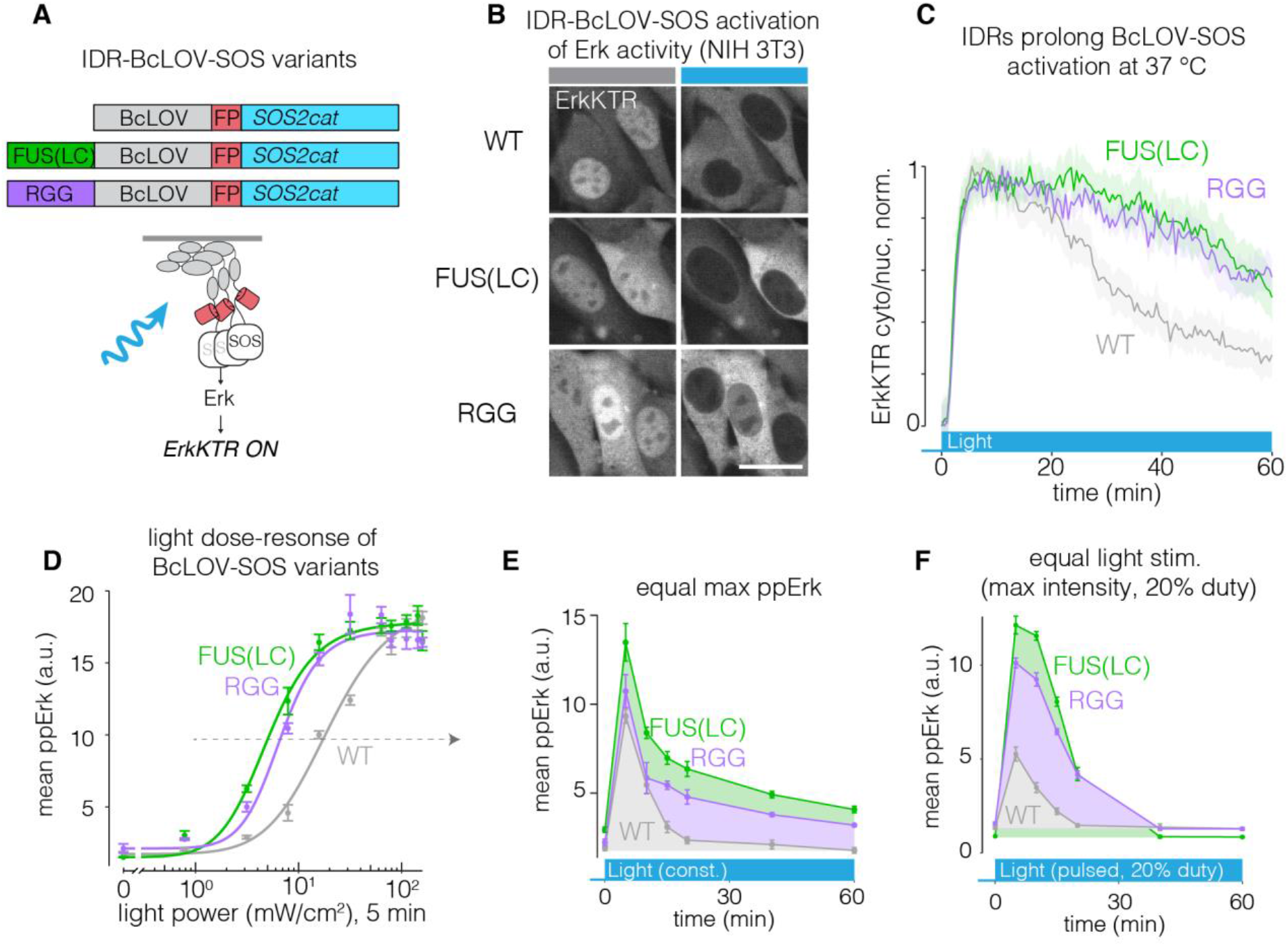
IDRs enhance sensitivity and strength of BcLOV-SOS_cat_. **A)** IDR-fused variants of BcLOV-SOS_cat_. **B)** The Ras/Erk pathway was activated in cells by IDR-BcLOV-SOS_cat_ variants in NIH 3T3 fibroblasts, as measured by the ErkKTR reporter. Scale bar = 20 μm. **C)** Quantification of ErkKTR activity during 1 hr of stimulation by BcLOV-SOS_cat_ variants. IDR variants show slower pathway decay. See **Supplementary Table 1** for details of optogenetic illumination parameters. **D)** Dose-response of light intensity on ppErk after 5 min of constant illumination at the indicated light dose. Data represent mean ± SEM of four replicates, each representing the mean signal from ~200-1000 cells. **E)** Comparison of ppErk activation dynamics by BcLOV-SOS_cat_ variants, each stimulated at a constant light intensity that produced equivalent max ppErk, as determined in **(D)** (dotted arrow). IDR variants showed higher sustained and integrated signaling over 1 hr of stimulation. Data represent the mean ± SEM of four replicates, each representing the mean of ~50-400 single cells. **F)** Comparison of ppErk activation dynamics by BcLOV-SOS_cat_ variants in response to pulsatile (20% duty cycle) maximum intensity light. IDR variants achieved > 2-fold higher maximal signal and more sustained and integrated activity compared to wt BcLOV-SOS_cat_. Data represent mean ± SEM of four replicates, each representing the mean of ~100-600 cells.

Taken together, our results for both BcLOV-EGFR and BcLOV-SOS_cat_ show that cluster strength serves as a tuning knob to that can offer stronger, more stable, and less perturbative stimulation of BcLOV-based optogenetic tools.

## Discussion

Our work shows that, upon light stimulation, the BcLOV4 protein both clusters and translocates to the plasma membrane, and that dual translocation and clustering can be leveraged for new optogenetic signaling probes including of the entire ErbB RTK family. Moreover, in contrast to previous evidence that clustering and translocation were mutually exclusive, we show that clustering promotes translocation of stimulated BcLOV4. Potentiation of clustering allowed for a higher amplitude of signal induction, increased sensitivity to light, extended durations of signaling, and a higher efficiency of signaling (signal per unit of BcLOV probe). BcLOV4 represents, to our knowledge, the second described photosensor whose light-induced clustering can be coopted for optogenetic control. The first, *Arabidopsis* Cry2^9^, has found widespread use across diverse systems of study^9,21,23–25,27,28,43^. As the importance of protein condensation continues to emerge^44^, we expect that BcLOV clustering will find extensive use cases, particularly for classes of condensation that occur at the membrane. Furthermore, the availability of multiple optogenetic clustering systems now provides more options for optogenetic control with distinct clustering properties (e.g. sensitivity, size, subcellular localization), and could further allow for multiplexed control of distinct clustering phenomena using the same blue light input.

We leveraged the dual translocation and clustering of BcLOV to regulate RTK signaling, with a focus on EGFR and the entire ErbB receptor family. These studies further demonstrate the remarkable modularity of BcLOV as an optogenetic actuator, building on its previous application to control GTPases, guanine nucleotide exchange factors (GEFs), GTPase activating proteins (GAPs), and phosphatidyl inositol-3 kinase (PI3K)^14–16^. We also found that by simply exchanging the intracellular domain of EGFR for the analogous domain of other ErbB family members, we could generate probes to control those receptors with no further optimization. Although this strategy also allowed stimulation of other families of RTKs including FGFR and PDGFR, we did observe RTK-family-specific effects including high basal signaling and limited activation strength. These results confirm the unique character of distinct RTK families that demands further optimization for their optimal activation, as has been observed previously^28,29,35,36^. These previous studies, as well as studies that optimized BcLOV for other signaling applications^14,15,26^, will provide a roadmap for future engineering of BcLOV-based RTK stimulation.

Our ability to enhance the strength and sensitivity of BcLOV through addition of disordered domains has important practical implications. Two potential complications of optogenetic approaches are 1) toxicity from extensive blue light stimulation, and 2) elevated basal levels of signaling from expression of the optogenetic probe. Potentiation of BcLOV membrane translocation (here using IDRs) addresses both of these concerns, allowing comparable signal induction with ~4-fold less light (**Figure 6D**), or with lower expression levels of the probe (**Figure S8**). In addition, and specifically for BcLOV, the increased sensitivity allows one to slow the spontaneous signal decay observed at mammalian temperatures (37 °C) in two ways. First, because decay depends on both light and temperature, lower light levels lead to slower decay^16^. Second, because IDRs can boost the amplitude of signal, the signal will remain above a given threshold for longer than for probes that lack the IDR.

While we found that IDR-induced enhancements in BcLOV translocation generally translated to signal activation of BcLOV-base optogenetic probes, we also found important exceptions. For example, despite increased membrane translocation arising from both the FUS(LC) and RGG IDRs (**Figure 4B,C**), only FUS-LC potentiated signaling of BcLOV-EGFR (**Figure 5D**). By contrast, both IDRs potentiated signaling of BcLOV-SOS_cat_ (**Figure 6D**). Furthermore, despite strong membrane localization over 2 hours with IDR-fused variants of BcLOV-mCh (**Figure 4C**), BcLOV-SOS_cat_ activity could only be extended, but not sustained indefinitely (**Figure 6F**). Future work will define the molecular details of BcLOV4 thermal sensitivity and provide additional strategies by which to mitigate or eliminate its effects. Collectively, these results highlight that a probe’s activation dynamics can be influenced by many factors, including the molecular nature of the signaling event, probe expression level, and the host cell environment. We also found that kinase activity of the probe can suppress its membrane translocation, and that such effects play a role in other widely used optogenetic systems as well (**Figure 3F, S7**).

In summary, BcLOV4 is a multifunctional photoreceptor that uniquely both clusters and translocates to the membrane in mammalian cells. BcLOV clustering can not only be leveraged for new types of single-component optogenetic tools, but can also be harnessed to enhance existing ones.

## Acknowledgements

The authors thank Erin Berlew for helpful discussions on this work. This work was supported by the National Institutes of Health (R35GM138211 for L.J.B) and the National Science Foundation (Graduate Research Fellowship Program to W.B., CAREER 2145699 to L.J.B., and CAREER MCB1652003 for B.Y.C). Cell sorting was performed on a BD FACSAria Fusion that was obtained through NIH S10 1S10OD026986 and is operated through the Penn Cytomics and Cell Sorting Resource Laboratory.

## Author Contributions

A.A.P., W.B., B.Y.C, and L.J.B. conceived the study. A.A.P, W.B., T.R.M. and L.J.B. performed experiments and analyzed data. L.J.B supervised the work. A.A.P. and L.J.B. wrote the manuscript and made figures, with editing from all authors.

---

**Supplementary Table 1.** Illumination and culture conditions for all experiments

| **Illumination and culture conditions for all experiments** |  |  |  |  |  |  |  |
| --- | --- | --- | --- | --- | --- | --- | --- |
| Figure | Temp (°C) | Duty Cycle | Intensity | ON time | OFF time | Humidified | CO<sub>2</sub> |
| 1B | 37 | 3.3% | 1.45W/cm<sup>2</sup> | 1s | 30s | Yes | 5% |
| 1D | 37 | 3.3% | 1.45W/cm<sup>2</sup> | 1s | 30s | Yes | 5% |
| 2C | 37 | 3.3% | 1.45W/cm<sup>2</sup> | 1s | 30s | Yes | 5% |
| 2D | 37 | 3.3% | 1.45W/cm<sup>2</sup> | 1s | 30s | Yes | 5% |
| 2E | 25,37 | 10% | 160mW/cm<sup>2</sup> | 0.5s | 5s | Yes | 5% |
| 2F-G | 37 | Variable | 160mW/cm<sup>2</sup> | 0.5s | Variable | Yes | 5% |
| 3B | 37 | 3.3% | 1.45W/cm<sup>2</sup> | 1s | 30s | Yes | 5% |
| 3C | 37 | 2.5% | 160mW/cm<sup>2</sup> | 0.5s | 20s | Yes | 5% |
| 3D<br>(5m ON) | 37 | 2.5% | 160mW/cm<sup>2</sup> | 0.5s | 20s | Yes | 5% |
| 3E | 37 | 3.3% | 1.45W/cm<sup>2</sup> | 1s | 30s | Yes | 5% |
| 3F | 37 | 3.3% | 1.45W/cm<sup>2</sup> | 1s | 30s | Yes | 5% |
| 4B | 37 | 3.3% | 1.45W/cm<sup>2</sup> | 1s | 30s | Yes | 5% |
| 4C | 37 | 3.3% | 1.45W/cm<sup>2</sup> | 1s | 30s | Yes | 5% |
| 5B | 37 | 3.3% | 1.45W/cm<sup>2</sup> | 1s | 30s | Yes | 5% |
| 5C | 37 | 100% | Variable | const. | ---- | Yes | 5% |
| 5D-E | 37 | Variable | 160mW/cm<sup>2</sup> | 0.5s | Variable | Yes | 5% |
| 5F | 37 | 2.5% | 160mW/cm<sup>2</sup> | 0.5s | 20s | Yes | 5% |
| 6B | 37 | 3.3% | 1.45W/cm<sup>2</sup> | 1s | 30s | Yes | 5% |
| 6C | 37 | 3.3% | 1.45W/cm<sup>2</sup> | 1s | 30s | Yes | 5% |
| 6D | 37 | 100% | Variable | const. | ---- | Yes | 5% |
| 6E | 37 | 100% | Variable | const. | ---- | Yes | 5% |
| 6F | 37 | 20% | 160mW/cm<sup>2</sup> | 0.5s | 2.5s | Yes | 5% |
| S1 | 37 | 3.3% | 1.45W/cm<sup>2</sup> | 1s | 30s | Yes | 5% |
| S2 | 37 | 5% | 160mW/cm<sup>2</sup> | 0.5s | 10s | Yes | 5% |
| S3-C | 37 | 3.3% | 1.45W/cm<sup>2</sup> | 1s | 30s | Yes | 5% |
| S3-E | 37 | 5% | 160mW/cm<sup>2</sup> | 0.5s | 10s | Yes | 5% |
| S4-B | 37 | 2.5% | 160mW/cm<sup>2</sup> | 0.5s | 20s | Yes | 5% |
| S5 | 37 | 3.3% | 1.45W/cm<sup>2</sup> | 1s | 30s | Yes | 5% |
| S6 (Live-cell Imaging) | 37 | 3.3% | 1.45W/cm<sup>2</sup> | 1s | 30s | Yes | 5% |
| S6 (IF Assay) | 37 | 2.5% | 160mW/cm<sup>2</sup> | 0.5s | 20s | Yes | 5% |
| S7 | 37 | 3.3% | 1.45W/cm<sup>2</sup> | 1s | 30s | Yes | 5% |
| SM1 | 37 | 3.3% | 1.45W/cm<sup>2</sup> | 1s | 30s | Yes | 5% |
| SM2 | 37 | 3.3% | 1.45W/cm<sup>2</sup> | 1s | 30s | Yes | 5% |
| SM3 | 37 | 3.3% | 1.45W/cm<sup>2</sup> | 1s | 30s | Yes | 5% |
| SM4 | 37 | 3.3% | 1.45W/cm<sup>2</sup> | 1s | 30s | Yes | 5% |
| SM5 | 37 | 3.3% | 1.45W/cm<sup>2</sup> | 1s | 30s | Yes | 5% |

**Supplementary Figure 1:**
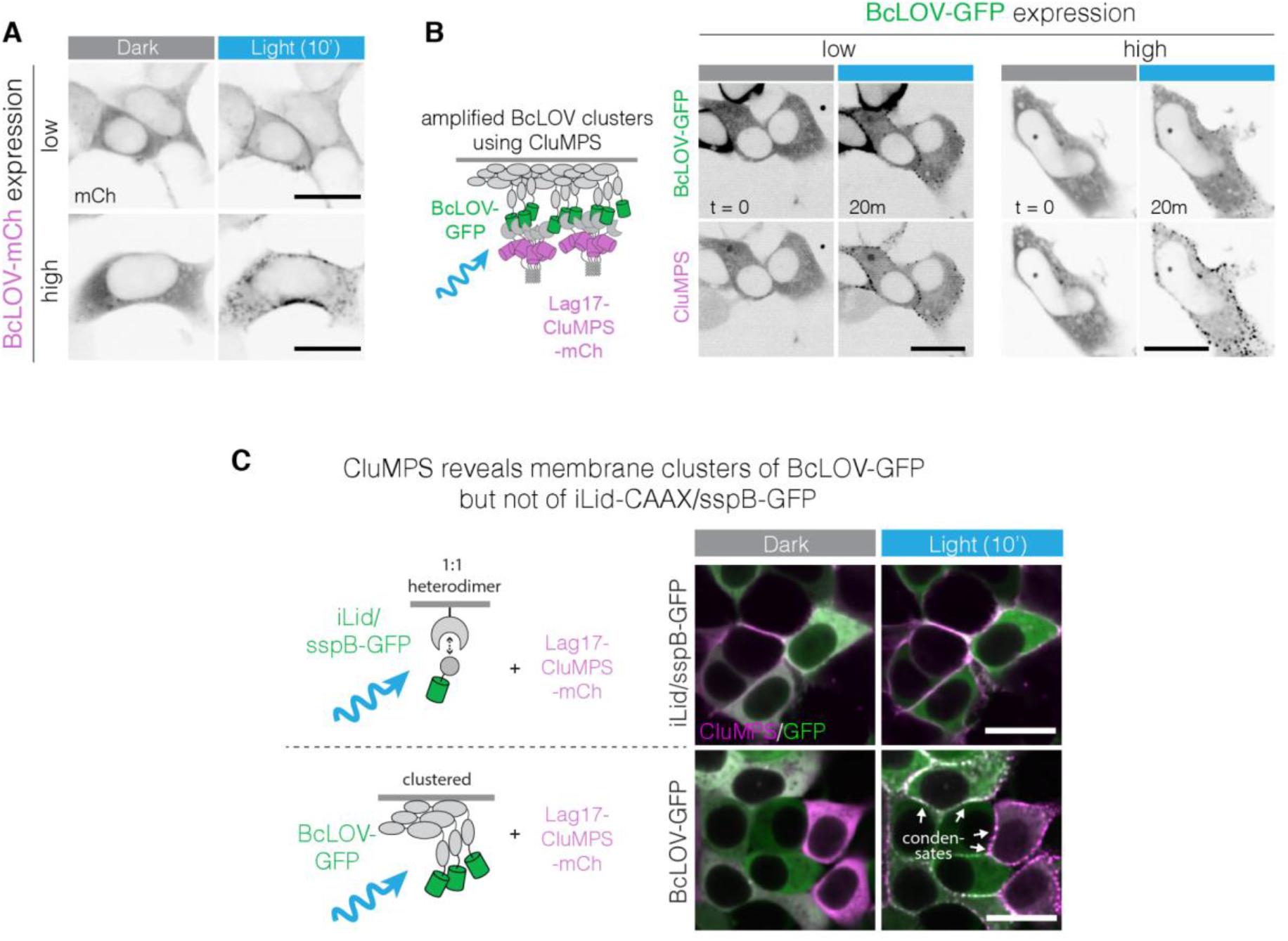
Light-induced clustering of BcLOV4 at the plasma membrane. **A)** Representative images of HEK 293T cells transiently transfected with BcLOV-mCherry under 40X confocal microscopy. In cells with high expression of BcLOV-mCherry, light-induced aggregation at the plasma membrane could be observed. Scale bar = 20 μm. **B)** Representative images of cells co-expressing BcLOV-GFP with a CluMPS reporter of GFP clustering (LaG17-CluMPS). Light-activated BcLOV demonstrated both membrane-association and condensate formation even with low levels of BcLOV4 expression. Scale bar = 20 μm. **C)** Representative images of membrane association and clustering of GFP when recruited by either iLid/sspB (1:1 heterodimer) or BcLOV4 in the presence of the CluMPS reporter. Light-induced recruitment through BcLOV4 resulted in membrane-associated GFP condensate formation, whereas recruitment through heterodimerization did not, suggesting that clustering and CluMPS activation is not simply a result of induced membrane translocation. Scale bars = 20 μm

**Supplementary Figure 2:**
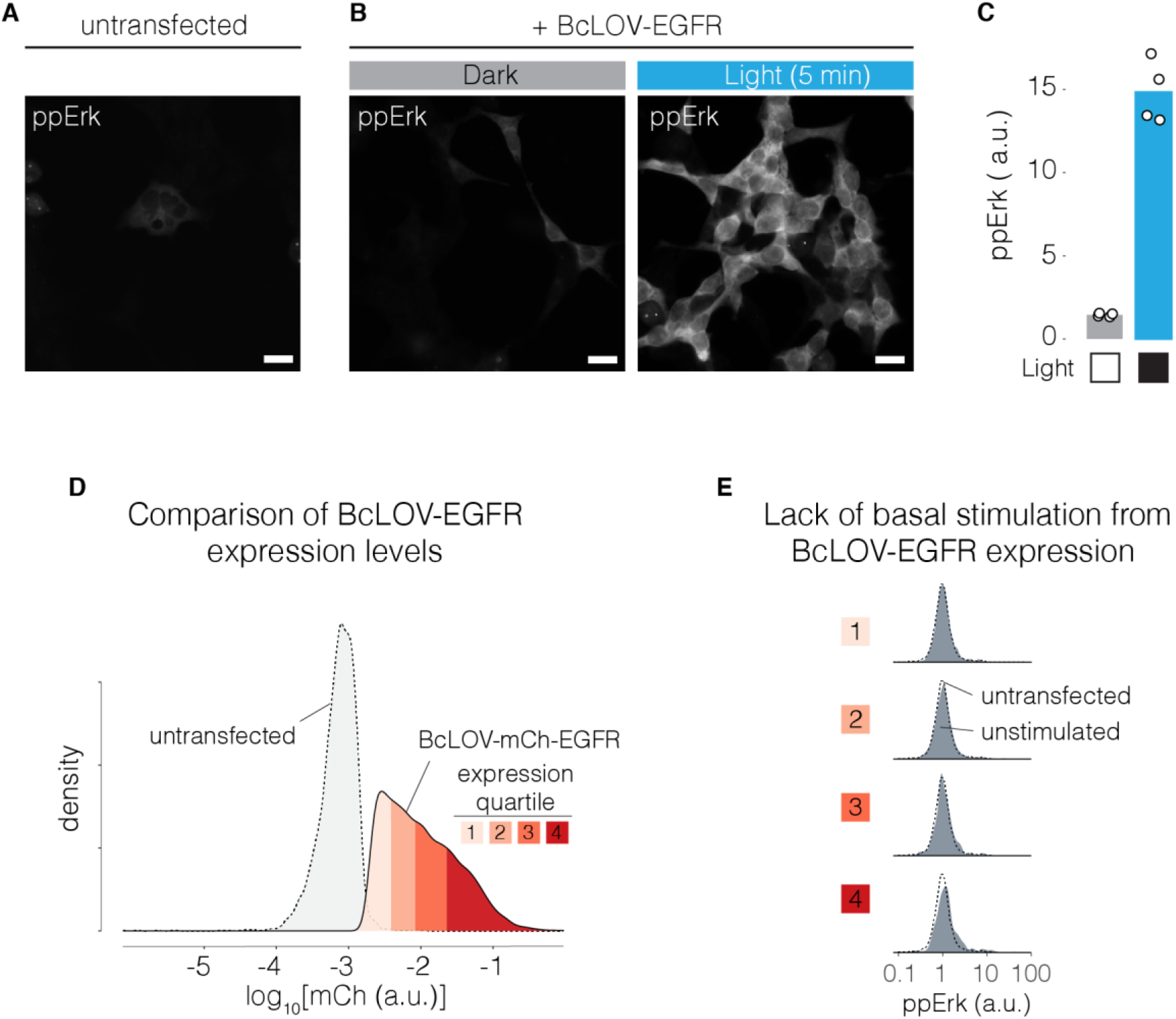
Optogenetic stimulation of Ras-Erk signaling using BcLOV-EGFR. Representative images of ppErk signal from untransfected HEK 293T cells (**A**) as well as from cells transfected with BcLOV-EGFR and kept in the dark or illuminated with 5 min of blue light (**B**). Scale bars = 20 μm. **C)** Quantification of ppErk immunofluorescence resulting from 5 min of optogenetic stimulation of BcLOV-EGFR. Data represent mean ppErk intensity from four replicates. Each replicate represents the mean of ~100 cells. See **Supplementary Table 1** for details of optogenetic stimulation. **D)** Visualization of expression level bins of BcLOV-EGFR vs untransfected HEK 293Ts. **E)** Single cell distributions of Erk activity in expressing (dark-state) vs non-expressing HEK 293T cells. Plots show no elevation of basal signaling resulting from probe expression.

**Supplementary Figure 3:**
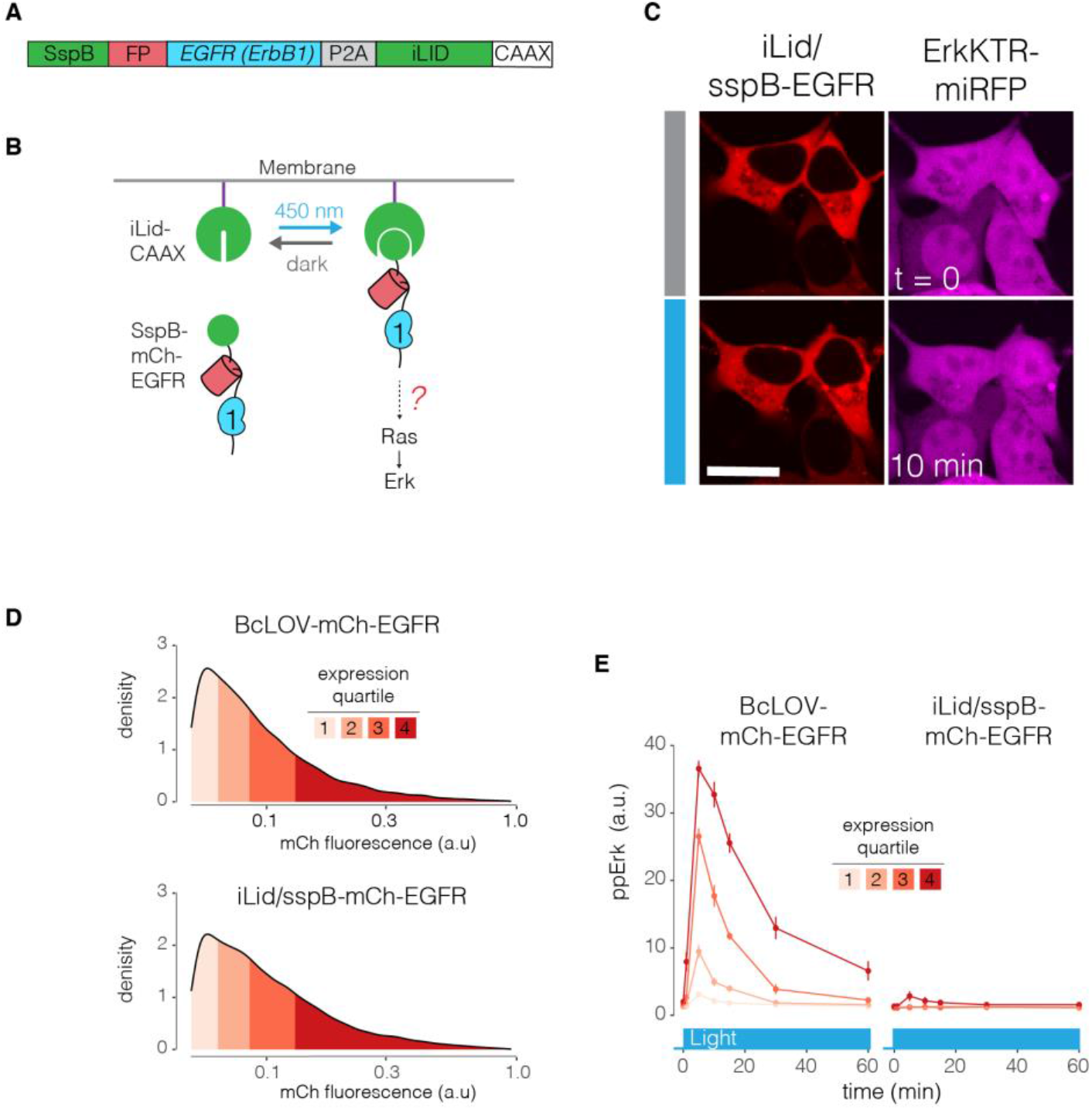
Membrane recruitment of the EGFR intracellular domain is not sufficient to activate downstream signaling. **A)** Schematic of iLID-CAAX/sspB-EGFR fusion construct. **B)** We asked whether light-induced recruitment of the EGFR intracellular domain to the plasma membrane was sufficient to activate downstream Ras-Erk signaling. **C)** Representative images of sspB-EGFR and ErkKTR-miRFP in HEK 293T cells in the presence and absence of light. Erk activation was not observed upon light stimulation. Scale bar = 20 μm. **D)** Single-cell mean expression levels of BcLOV- or iLid/sspB-based EGFR tools. Colors represent expression quartiles. **E)** Immunofluorescence of ppErk levels downstream of either BcLOV-EGFR or iLid/sspB-EGFR after light stimulation, separated by expression level. Expression-level-dependent stimulation was observed for BcLOV-EGFR. By comparison, no signaling was observed for iLid/sspB-EGFR except for a small amount at the very highest expression levels. Data represents the mean ± SEM of four replicates, with each replicate representing the mean of ~10-120 cells per expression quartile. See **Supplementary Table 1** for details of optogenetic stimulation parameters.

**Supplementary Figure 4:**
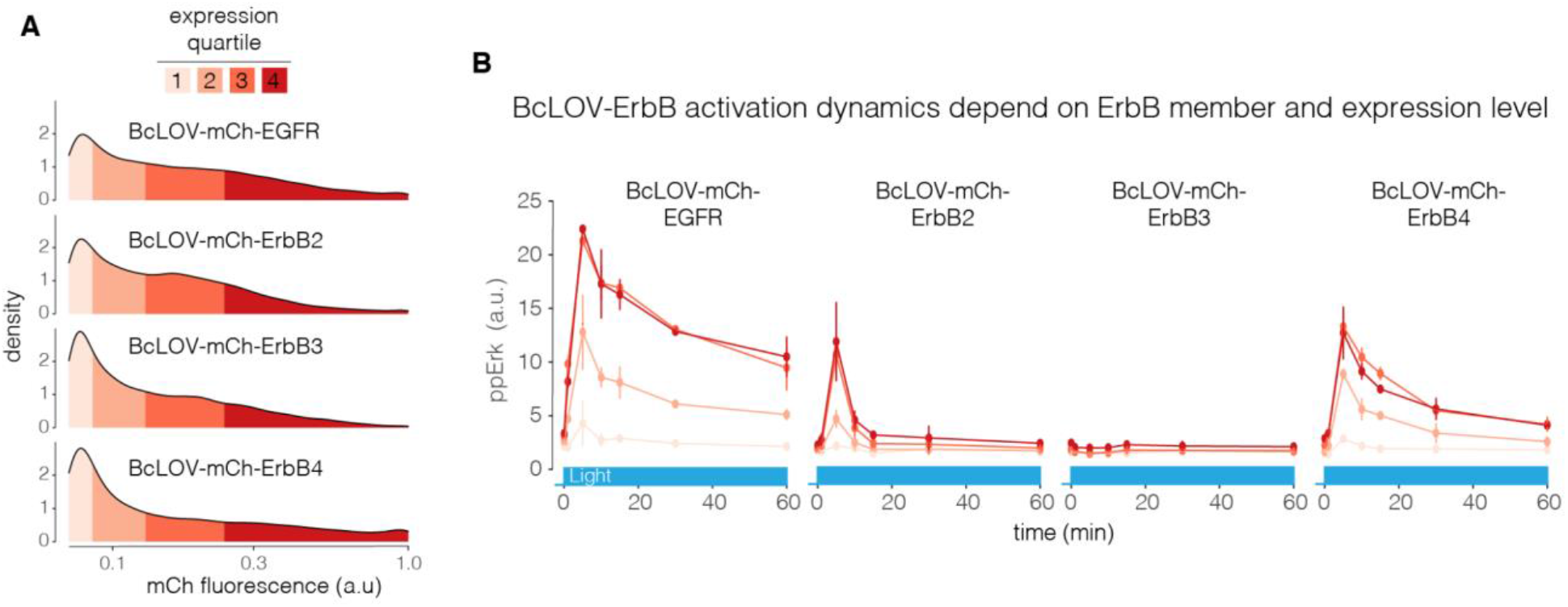
ppErk dynamics downstream of BcLOV-ErbB stimulation depend on the ErbB family member and on the expression level of the probe. **A)** Single-cell mean expression levels of BcLOV- or iLid/sspB-based EGFR tools. Colors represent expression quartiles. **B)** Comparison of differential magnitude, dynamics, and duration of ppErk signaling downstream of each ErbB member, visualized as a function of expression level. Data represent the mean ± SEM of two replicates, with each replicate representing the mean of ~200-800 cells per expression quartile. See **Supplementary Table 1** for optogenetic stimulation parameters.

**Supplementary Figure 5:**
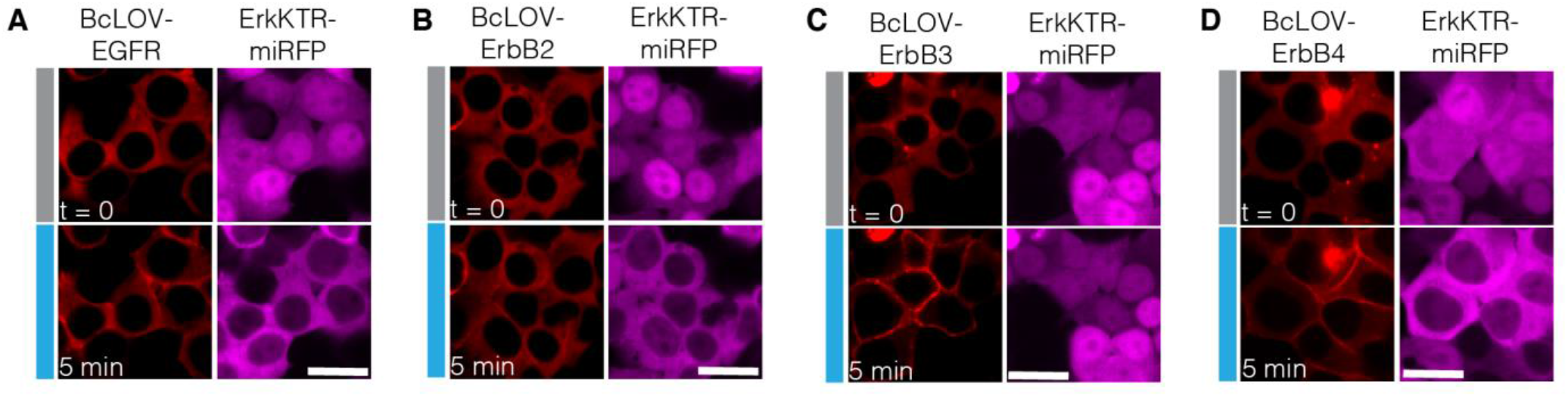
ErkKTR stimulation by BcLOV-ErbB1-4. Representative images of the ErkKTR-miRFP reporter in HEK 293T cells that were transiently transfected with BcLOV-ErbB1-4. Scale bars = 20 μm. See **Supplementary Table 1** for optogenetic stimulation parameters.

**Supplementary Figure 6:**
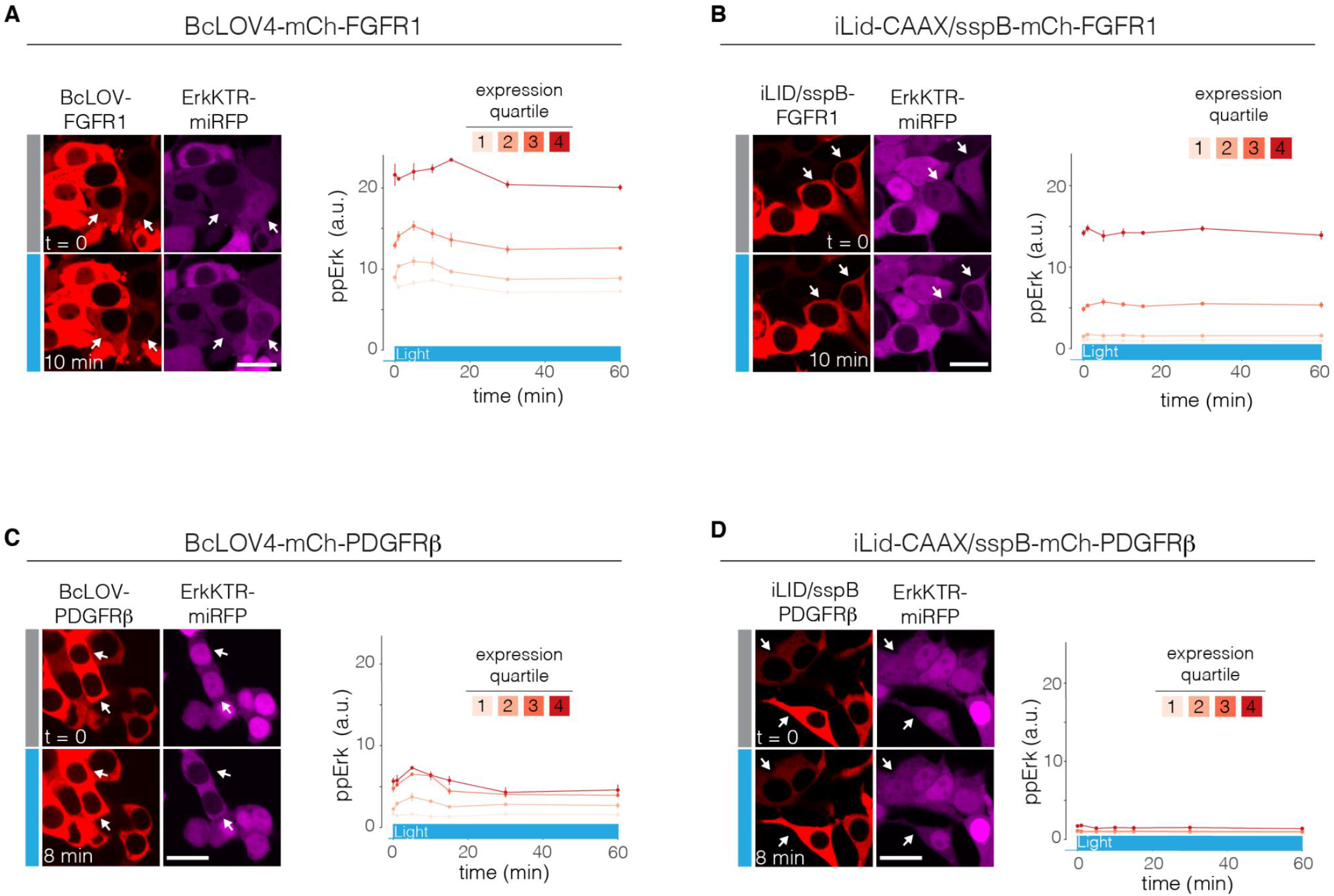
Stimulation of FGFR1 and PDGFRβ with BcLOV4 and comparison with stimulation through 1:1 heterodimeric recruitment to the plasma membrane. **A)** Stimulation of FGFR1 signaling with BcLOV4. Representative images (left) of the ErkKTR-miRFP reporter in HEK 293T cells that were transiently transfected with BcLOV-FGFR1. Light-induced ErkKTR stimulation can be observed only in low-expressing cells (white arrows). High-expressing cells show constitutively high Erk activity even in the absence of light. Immunofluorescence for ppErk (right) confirms expression-level dependence of basal levels and light-induced signaling. **B)** Stimulation of FGFR signaling through membrane recruitment with 1:1 heterodimerization of iLid/sspB. Representative images (left) of ErkKTR activity in HEK 293T cells that express iLid-CAAX/sspB-FGFR1. Here, membrane recruitment can stimulate the pathway at intermediate-high expression levels (white arrows). As in (**A**), high-expressing cells show constitutive activity even in the dark. Immunofluorescence (right) confirms basal Erk activity for cells at high expression levels. **C)** Stimulation of PDGFRβ signaling with BcLOV4. Representative images (left) of the ErkKTR-miRFP reporter in HEK 293T cells that were transiently transfected with BcLOV-PDGFRβ. Light-induced ErkKTR stimulation can be observed only in mid-high-expressing cells (white arrows). High-expressing cells show constitutively high Erk activity even in the absence of light. Immunofluorescence for ppErk (right) shows expression-level dependence on Erk phosphorylation. **D)** Stimulation of PDGFRβ signaling through membrane recruitment with 1:1 heterodimerization of iLid/sspB. Representative images (left) of ErkKTR activity in HEK 293T cells that express iLid-CAAX/sspB-PDGFRβ. Unlike for FGFR1, membrane recruitment does not stimulate PDGFRβ and downstream Erk activity (white arrows). Scale bars = 20 μm. Immunofluorescence (right) confirms no Erk activation across all expression levels. For immunofluorescence, data represents the mean ± SEM of two replicates, with each replicate representing the mean of ~100-600 cells per expression quartile. See **Supplementary Table 1** for details of optogenetic illumination parameters.

**Supplementary Figure 7:**
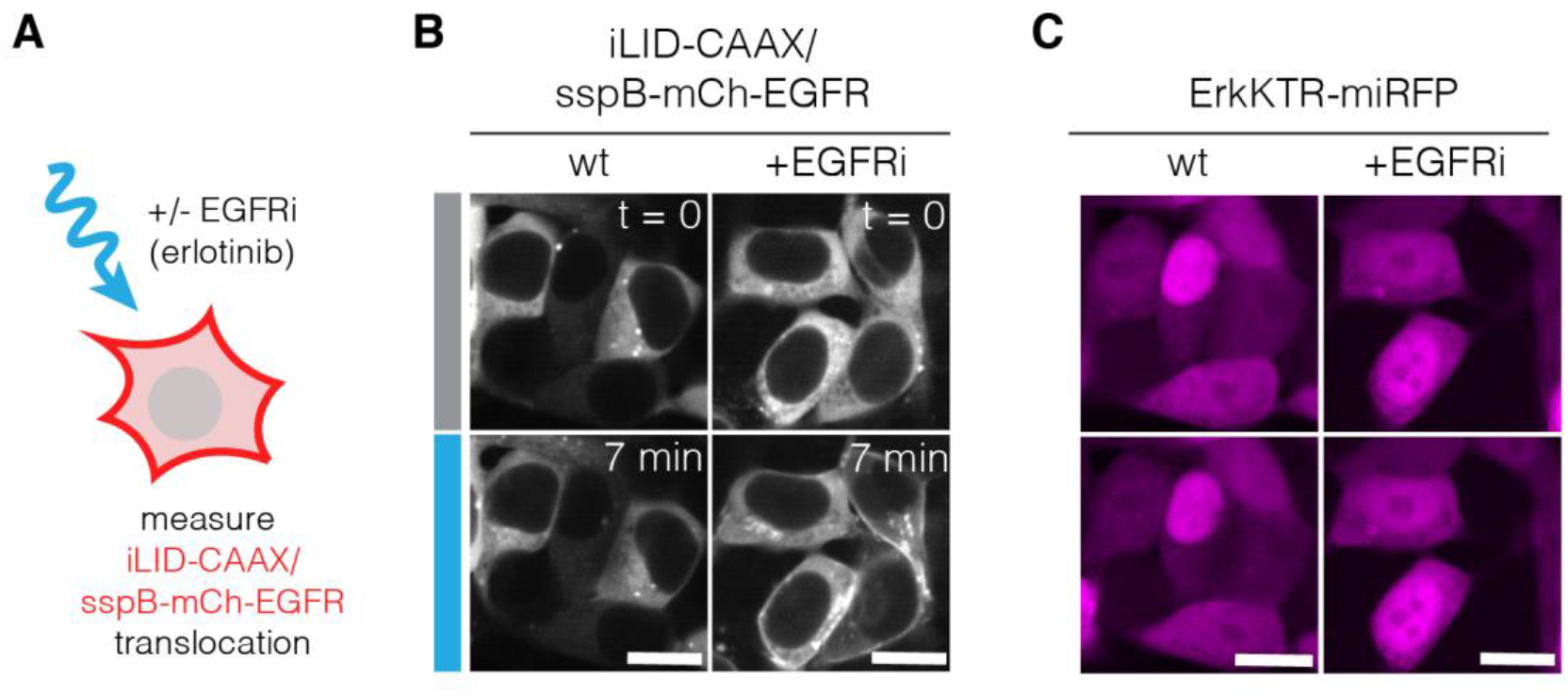
EGFR kinase activity suppresses iLID/SspB-mediated membrane translocation. **A)** HEK 293T cells expressing iLID-CAAX/SspB-mCh-EGFR were treated with 1uM of erlotinib (EGFR inhibitor) prior to blue-light stimulation, and membrane translocation and ErkKTR activity was observed for wt and drug-treated conditions. **B)** In wt cells, minimal membrane translocation could be observed. However, in cells treated with erlotinib, clear translocation was observed in response to light stimulation. **C)** Corresponding ErkKTR-miRFP images for cells in (**B**). ErkKTR activity could not be observed in either the presence or absence of EGFRi, regardless of sspB-EGFR recruitment, due to inhibition of the EGFR kinase. Scale bars = 20 μm. See **Supplementary Table 1** for details of optogenetic illumination parameters.

## Supplementary Movie Captions

**Supplementary Movie 1. CluMPS amplifies and visualizes membrane-associated BcLOV4 condensates.** Confocal imaging of BcLOV-GFP translocation with or without CluMPS in HEK 293T cells upon stimulation with 488 nm light. Time is in minutes:seconds. Blue square indicates light stimulation. Scale bar = 10 μm.

**Supplementary Movie 2. The Ras/Erk pathway was activated in cells by BcLOV-EGFR.** Confocal imaging of BcLOV-EGFR translocation and activation of ErkKTR-miRFP in HEK 293T cells upon stimulation with 488 nm light. Blue square indicates light stimulation.

**Supplementary Movie 3. Magnitude of membrane translocation differs between fusions of ErbB family members with BcLOV4.** Confocal imaging of BcLOV-mCh-ErbB1-4 translocation to the membrane upon blue light stimulation with 488 nm light. Blue square indicates light stimulation.

**Supplementary Movie 4. ErbB activity correlates with membrane ruffling.** Confocal imaging of BcLOV-mCh-ErbB1-4 translocation and induced membrane ruffling upon stimulation with 488 nm light. Blue square indicates light stimulation.

**Supplementary Movie 5. Disordered domains increase light-induced membrane translocation of BcLOV4.** Confocal imaging of IDR-fused variants of BcLOV-mCh in HEK 293T cells upon stimulation with 488 nm of light. Blue square indicates light stimulation.

## Methods

### Cell culture

All cell lines were maintained at 37°C and 5% CO_2_ in a cell culture incubator. Lenti-X HEK 293T cells were cultured in DMEM containing 10% fetal bovine serum (FBS) and 1% penicillin/streptomycin (P/S). NIH 3T3 fibroblast cells were cultured in DMEM containing 10% calf serum (CS) and 1% P/S.

### Plasmid design and assembly

Constructs were assembled using Gibson assembly. DNA fragments for the inserts and backbone were generated via PCR with primers obtained from Genewiz (Azenta Life Sciences), and inserted into the backbone using HiFi cloning mix (New England Biolabs). All constructs were verified with Sanger sequencing. DNA sequences encoding BcLOV4 was a kind gift from Dr. Brian Y. Chow^11^. GFP-binding nanobody LaG17 was obtained from Dr. Michael P. Rout^45^. LaG17-CluMPS was previously described^17^. ErkKTR-miRFP670 was previously described^16^. EGFR/ErbB1 was sourced from Opto-hEGFR, which was a kind gift from Dr. Harold Janovjak^35^. ErbB2 was sourced from MSCV-human Erbb2-IRES-GFP, which was a gift from Martine Roussel (Addgene plasmid # 91888; http://n2t.net/addgene:91888; RRID:Addgene_91888). ErbB3 was sourced from pDONR223-ERBB3, which was a gift from William Hahn & David Root (Addgene plasmid # 23874; http://n2t.net/addgene:23874; RRID:Addgene_23874). ErbB4 was sourced from pDONR223-ERBB4, which was a gift from William Hahn & David Root (Addgene plasmid # 23875; http://n2t.net/addgene:23875; RRID:Addgene_23875). iLID, sspB, and SOS_cat_ were sourced from previously described constructs^16^. FUS(LC) (1-163) and LAF-1 RGG were kindly provided by Dr. Matthew Good.

### Plasmid transfection

Lenti-X HEK 293T cells were transfected using Lipofectamine™ 3000 transfection reagent (ThermoFisher Scientific) following the manufacturer’s protocol. The transfection mixture contained 100 ng/μl DNA, 2% Lipofectamine™ reagent and 2% P3000 reagent, and was brought up to a final volume of 10μl with Opti-MEM™ (ThermoFisher Scientific). The transfection mixture was incubated for 15 minutes at room temperature and was then added to the cells. For cells seeded in 96-well plates, 10 μl of transfection mixture was added per well. Cells seeded in 384-well plates received 2.5 μl of transfection mixture per well.

### Lentiviral packaging and cell line generation

Lentivirus was packaged by contransfecting the pHR transfer vector, pCMV-dR8.91 (Addgene, Catalog #12263), and pMD2.G (Addgene, Catalog #12259) into Lenti-X HEK 293T cells. Cells were seeded one day prior to transfections at a density of 700,000 cells/mL in a six-well plate. Cells were transfected using the calcium phosphate transfection method: for 300μl of transfection mix, 1.5μg of transfer vector, 1.33μg of pCMV-dR8.91, 0.17μg of pMD2.G, 150μl of 2X HEPES-buffered saline (HeBS) and H_2_O up to 132ul were mixed. Then, 18 μl of 2.5 mM CaCl2 was then added, the mixture was incubated for 1 minute 45 seconds at room temperature, and then the mixture was added dropwise to the cells. One day post-transfection, media was removed from the plate and replaced with fresh media. Two days post-transfection, media containing virus was collected and centrifuged at 800 x g for 3 min. Supernatant from centrifuged media was then collected and filtered through a 0.45-μm filter. 500 μL of filtered virus was added to 100,000 cells (Lenti-X HEK293T or NIH 3T3) seeded in a six-well plate. Cells were observed for transduction by checking for fluorescence ~1-2 days post-infection. Cells were expanded over multiple passages. Successfully transduced cells were enriched through cell sorting using a BD FACSAria Fusion.

### Preparation of cells for plate-based experiments

For experiments, cells were seeded in 96- or 384-well plates (Cellvis 96-well plate with glass-like polymer bottom, catalog number P96-1.5P; Greiner Bio-One CELLSTAR 384-well, Cell Culture-Treated, Flat-Bottom Microplate, catalog number 781091). First, wells were coated with 30 μL (for 96-well plate) or 12 μL (for 384- well plate) of 10 μg/ml of MilliporeSigma™ 597 Chemicon™ Human Plasma Fibronectin Purified Protein in 1X PBS for 15 minutes at 37°C. For 96-well plate experiments, 25,000 Lenti X HEK 293T or 12,000 NIH 3T3 cells were seeded in 150 μl of P/S-free cell-culture medium (DMEM + 10% FBS or 10% CS) in each well. For 384-well plate experiments, 3500 Lenti-X HEK 293T or 2500 NIH 3T3’s were seeded. Following the seeding step, the plates were spun down at 20 x g for 1 minute to promote uniform distribution of cells throughout the well.

For experiments with stable cell lines, cells were starved after 24 hours by performing seven 80% washes (for 384-well plate) or four 75% washes (for 96-well plate) with starvation media (DMEM +1% P/S) using an automated plate washer (BioTEK ELx405). Experiments were performed 3-4 hours post-starvation. For experiments with transiently transfected cells, cells were starved 6 hours post-transfection to remove lipofectamine reagent from cells, and experiments were performed after overnight starvation.

### Optogenetic stimulation

For live-cell imaging experiments, the 488 nm laser was used to stimulate BcLOV4 tools for membrane translocation. For fixed-cell experiments, cells were stimulated with a single-color blue LED optoPlate-96^42^. LED intensities were calibrated using a Thorlabs power meter (catalog number PM16-140). Briefly, each well of the optoPlate was turned on to maximum intensity. The power meter was used to scan the well, and the maximum intensity reading from that well was recorded. This process was repeated for all wells. The ratio of each LED intensity to the dimmest LED intensity found was then calculated, and this value was used as a “scaling factor”, such that each LED was scaled down to emit at the same intensity as the weakest LED. In this way, all LEDs were set to the same power output. For stimulation experiments, the light intensity was configured to stimulate the wells with a range of light intensities spanning from 0 to 160 mW/cm^2^. The Arduino IDE (v1.8) was used to program the Arduino Micro present on the optoPlate-96. A 20 mm tall black adaptor was used for even light diffusion across each of the wells on the 384-well plate. For time course experiments, time points were assigned to individual wells, and stimulations were run in a sequential manner to allow simultaneous fixing of cells at the end of the experiment. The apparatus was arranged inside a standard cell culture incubator set at 37°C and 5% CO2, and the experiments were run under dark conditions to avoid unwanted light exposure. Prior to experiments, optoPlate stimulation protocols were tested to ensure that no sample heating occurred due to heat generation from the device. Sample temperatures were measured using a custom-built immersion temperature sensor.

### Immunofluorescence staining

Immediately following completion of a stimulation protocol,16% paraformaldehyde (Paraformaldehyde Aqueous Solution, Electron Microscopy Sciences, catalog number 15710) was added to each well to a final concentration of 4%, and cells were incubated for 10 minutes in the dark. Cells were then permeabilized 1X PBS + 0.1% Triton X-100 for 10 minutes at room temperature (RT). Cells were further permeabilized with icecold 100% methanol at −20°C for 10 minutes. After permeabilization, cells were blocked with 1% BSA in 1X PBS for 30 minutes at RT. Cells were then incubated in primary antibody diluted in 1X PBS + 0.1% BSA (phospho-p44/42 MAPK (Erk1/2) (Thr202/Tyr204), Cell Signaling, catalog number 4370L, 1:400 dilution) at 4°C overnight. After overnight incubation, the primary antibody was removed and the plate was washed 5 times in PBS + 0.1% Tween-20 (PBS-T). Cells were then incubated with secondary antibody (Jackson Immunoresearch Alexa Fluor 488 AffiniPure goat anti-rabbit IgG (H+L), 1:500) and 4,6-diamidino-2-phenylindole (DAPI; ThermoFisher Scientific, catalog number D1306, 300 nM) in 1X PBS + 0.1% BSA for 1 hour at RT. The secondary antibody was removed and the plate was washed 5 times in PBS-T.

### Imaging

#### Live-cell imaging

Live-cell imaging was done using a Nikon Ti2E microscope equipped with a Yokagawa CSU-W1 spinning disk, 405/488/561/640nm laser lines, an sCMOS camera (Photometrics), a motorized stage and an environmental chamber (Okolabs). HEK 293T and NIH 3T3 cells with desired constructs were plated in 96- or 384-well plates and imaged with a 40X oil immersion objective at 37°C and 5% CO_2_. For the EGFR inhibition experiments, cells were treated with 1 μM of erlotinib 30 minutes prior to imaging. Erlotinib was kindly provided by Dr. Arjun Raj.

#### High-content imaging

For fixed-cell experiments, samples were imaged using a Nikon Ti2E epifluorescence microscope equipped with DAPI/FITC/Texas Red/Cy5 filter cubes, a SOLA SEII 365 LED light source and motorized stage. High-content imaging was performed using the Nikon Elements AR software. Image focus was ensured using image-based focusing in the DAPI channel.

### Image processing and analysis

#### Live-cell ErkKTR quantification

To determine the cytoplasmic/nuclear fluorescence ratios of ErkKTR reporter from the live-cell imaging experiments for Figures 2D, ImageJ^46^ was used to manually compare the pixel intensities of the mean cytoplasmic and nuclear intensities for 25 cells in the same field of view. The obtained values were exported into R for data analysis using the dplyr^47^ and ggplot2^48^ packages.

#### Immunofluorescence quantification

Cell Profiler^49^ was used to quantify ppErk levels in the fixed-cell experiments. Cells were segmented using the DAPI channel and the cytoplasm was identified by expanding a 5-pixel ring from the nuclei. The obtained cytoplasmic and nuclear fluorescence values were exported into R for data analysis using the dplyr and ggplot2 packages.

#### Membrane recruitment

Membrane recruitment of BcLOV4 in Figure 4C was quantified using the MorphoLibJ Plugin for ImageJ^50^. All experiments were performed in cells stably expressing a fluorescent membrane marker (GFP-CAAX). Images of the membrane marker were used to automatically segment single cells using the “Morphological Segmentation” feature of the MophoLibJ with a threshold of 150. Segmentation of each membrane marker image was exported as a separate tiff image. Segmentation images were imported to CellProfiler along with the corresponding images of BcLOV-mCherry variants. Membrane values of mCh were then determined by designating a 1-pixel-wide perimeter of each cell’s membrane. The membrane intensity and total cell intensity of BcLOV4 was then measured and recorded for each cell. R was used to process these values, normalizing membrane BcLOV4 intensity of each cell by the whole cell intensity and averaging these single cell values for each time point.

#### Curve fitting

ppErk levels for the dose-response curves of IDR-fused variants of BcLOV-EGFR and BcLOV-SOS_cat_ (**Figure 5D and 6D**) were fit to a Hill function of the form

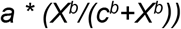

where *X* is the power of light used, *a* is the maximal amount of Erk activation, *b* is the parameter defining steepness of the curve and *c* is the percentage of light needed to achieve half-maximal activation of Erk. A MATLAB function was written to determine the parameters, and the curves were fitted on RStudio.

### Constructs used in this study

| Construct | Backbone + Promoter |
| --- | --- |
| BcLOV-GFP | pHR CMV |
| SspB-mCherry-2A-iLID-CAAX | pHR CMV |
| LaG17-mCh-HOTag3 | pHR PGK |
| ErkKTR-miRFP670 | pHR SFFV |
| BcLOV-mCherry-EGFR/ErbB1 | pHR CMV |
| BcLOV-mCherry-ErbB2 | pHR CMV |
| BcLOV-mCherry-ErbB3 | pHR CMV |
| BcLOV-mCherry-ErbB4 | pHR CMV |
| SspB-mCherry-EGFR-2A-iLID-CAAX | pHR CMV |
| BcLOV-mCherry-FGFR | pHR CMV |
| BcLOV-mCherry-PDGFRβ | pHR CMV |
| SspB-mCherry-FGFR-2A-iLID-CAAX | pHR CMV |
| SspB-mCherry-PDGFR-2A-iLID-CAAX | pHR CMV |
| BcLOV-mCherry | pcDNA3.1 CMV |
| FUS(LC)-BcLOV-mCherry | pcDNA3.1 CMV |
| RGG-BcLOV-mCherry | pcDNA3.1 CMV |
| BcLOV-mCherry-EGFR | pHR CMV |
| FUS(LC)-BcLOV-mCherry-EGFR | pHR CMV |
| RGG-BcLOV-mCherry-EGFR | pHR CMV |
| BcLOV-mCherry-SOS<sub>cat</sub> | pHR SFFV |
| FUS(LC)-BcLOV-mCherry-SOS<sub>cat</sub> | pHR SFFV |
| RGG-BcLOV-mCherry-SOS<sub>cat</sub> | pHR SFFV |
| GFP-CAAX | pHR CMV |

